# QuoVidi: a open-source web application for the organisation of large scale biological treasure hunts

**DOI:** 10.1101/2020.06.30.177006

**Authors:** Guillaume Lobet, Charlotte Descamps, Lola Leveau, Alain Guillet, Jean-François Rees

## Abstract

Learning biology, and in particular systematics, requires learning a substantial amount of specific vocabulary, both for botanical and zoological studies. While crucial, the precise identification of structures serving as evolutionary traits and systematic criteria is not per se a highly motivating task for students. Teaching this in a traditional teaching setting is quite challenging especially with a large crowd of students to be kept engaged. This is even more difficult if, as during the COVID-19 crisis, students are not allowed to access laboratories for hands-on observation on fresh specimens and sometimes restricted to short-range movements outside their home.

Here we present QuoVidi, a new open-source web platform for the organisation of large scale treasure hunts. The platform works as follows: students, organised in teams, receive a list of quests that contain morphologic, ecologic or systematic terms. They have to first understand the meaning of the quests, then go and find them in the environment. Once they find the organism corresponding to a quest, they upload a geotagged picture of their finding and submit this on the platform. The correctness of each submission is evaluated by the staff. During the COVID-19 lockdown, previously validated pictures were also submitted for evaluation to students that were locked in low-biodiversity areas. From a research perspective, the system enables the creation of large image databases by the students, similar to citizen-science projects.

Beside the enhanced motivation of students to learn the vocabulary and perform observations on self-found specimens, this system allows faculties to remotely follow and assess the work performed by large numbers of students. The interface is freely available, open-source and customizable. It can be used in other disciplines with adapted quests and we expect it to be of interest in many classroom settings.

## Introduction

Teaching biology to first-year bachelor students is a challenge. As educators, our aim is usually twofold. First, we want the students to learn a new set of knowledge and integrate it. Second, and this is for us equally important, we want the students to engage with the topic at hand. We want to transmit our passion and curiosity about the topic that we teach. Third, we also want students to learn to observe the world around them. It is one thing to learn a topic from a textbook, it is another to observe it in real life. However, the main issue is that the classroom is, often by design, completely disconnected from the natural world. The challenge is therefore to find a way for students to learn and engage with biology, despite that given disconnection. Last but not least, in the Spring semester of 2020 (January to June) it was necessary for us to adapt the learning activities to the containment measures related to COVID-19.

The formal aim of our biology course – given in the Bioengineering Faculty, UCLouvain, Belgium – is to discover plant and animal structures, organs and their function at the individual scale. To achieve this, students need to learn specific vocabulary related to these structures. The classic way to present this vocabulary to a student audience is to review a series of slides illustrating these different characteristics. This vocabulary is usually very boring for teachers to describe (imagine the slides showing all the different shapes of leaves) and the content is not very interesting for students to listen to either. Yet this vocabulary is an important prerequisite for describing any biological structure and for later systematic identification of taxons using dichotomous keys. Its learning is essential. The question is therefore how to make this learning process motivating for the students and give them the opportunity to learn over time instead of memorising a list of words? The additional difficulty is that this learning activity must be able to be set up with more than 300 students and few teaching resources.

To create this learning activity, we decided to draw inspiration from all the pedagogical techniques that aim to place the student at the centre of his learning. Student-centred learning and active learning emerged as important pedagogical techniques during the last century [REF]. Active learning is characterised by (i) involving the student in the construction of his or her learning, (ii) engaging the student in an in-depth treatment of the subject matter, (iii) constructing learning through interaction (with the teacher or other students), (iv) conceiving of learning as the evolution of knowledge and skills [1,2]. Studies have shown that the more cognitively and socially engaged the student is in a learning task, the more perennial the learning task becomes [1,3]. Active learning improves the performance of students and acts to reduce the gap achievement between advantaged and disadvantaged students [4]. In order to stimulate learning through interaction and create a collective emulation around this activity, the idea of creating a campus-wide biological treasure hunt finally emerged from the discussions. Beyond simply being active through the manipulation of information, the student has to transform and produce new information that is not provided in the learning material.

Gamification is another recent technique to better engage the students in a learning activity. Gamification is defined by [5] as “game-based mechanics, aesthetics, and game thinking to engage people, motivate action, promote learning, and solve problems”. In many studies, students’ levels of engagement increased significantly following the introduction of game elements, such as points, challenges, quests or progress bar [6]. The gamified environnement can afford intrinsic motivation and engagement, which are also targeted by active learning.

To assemble these different elements – biological vocabulary, observation, active learning and gamification – in a comprehensive learning activity, we created a large scale biological treasure hunt for our students. In short, we provided students with a list of specific biological vocabulary. They had to understand the list and find the different elements outside of the classroom, in the natural world. External resources (books, selected websites, wiki pages) describing this vocabulary were available to them. Complexity of understanding (some words are more difficult than others) as well as the difficulty of identification in the field were rewarded with different points.

To manage the treasure hunt, we designed a new web-based platform, QuoVidi (which would loosely translate from latin as “where did you see”), for the organisation of large scale, decentralised, biological treasure hunts. QuoVidi is an open-source project available at www.quovidi.xyz. The objective of this publication is to describe the project, to show how we were able to adapt this learning activity to the covid-19 crisis, and finally, to show the results and success of the activity with the students.

## Presentation of QuoVidi

QuoVidi is a web application for the organisation and management of large scale biological treasure hunts. It was created to teach students to learn new biological terms (both in zoology and botany) and to teach them to observe the natural world surrounding them.

### Setting up the activity

First, educators have to prepare a list of quests to find in the natural world.

These quests should be tailored and adapted for the target public. For instance, in our experience with first year biology students, the quests revolved around biological structures and families (tab. 1). Each quest is given a specific reward (points) depending on its intrinsic difficulty and rareness. Quests can be sorted in different groups (for instance “animal” and “plant”) and subgroups (for instance “animal species” and “leaf shapes”) to help students navigate them.

**Table 1:** Examples of quests used in the QuoVidi activity. Quests are sorted in the different groups and subgroups to help students navigate them. Each quest yields a number of points depending on its difficulty.

| quest | group | subgroup | points |
| --- | --- | --- | --- |
| Find an achene | plant | types of fruits | 1 |
| Find a flower with a bilateral symmetry | plant | types of flowers | 2 |
| Find a Siphonaptera | animal | animal groups | 3 |
| Find an example of aposematism | animal | animal physical attributes | 1 |

Second, educators have to assign students to groups to perform the activity. Students in the same group will be able to share pictures and collaborate on the data collection. When logging into the web interface, students will be able to see the collected pictures and rewards from their own group. They will also be able to see the total number of points of the competing groups.

Educators also have the possibility to define specific game parameters, such as specific geographic regions in which the game takes place or restriction on the number of submissions in each quest group (adding for instance a point penalty below a certain number of “animal” or “plant” submissions).

Once the list of quests, users and groups are defined, the activity can start. Two main activities are available for the students: an *in situ* treasure hunt and an *ex situ* photo quiz activity.

### Treasure hunt

The main activity of the platform is the biological treasure hunt. Students have to go outside (although some of the creatures may be also found in their home such as food parasites, e.g. Lepisma sp. or flies) to find the different quests setup by the educators. Once they find a specific quest, they have to take a picture of it with their smartphone. We ask the student to take unambiguous pictures, where the subject of the quest is clearly identified and visible. We also ask them to leave the natural environment intact, without killing any plant or animal in the process.

They can then store the picture on the QuoVidi web interface. When stored, pictures are automatically resized (for efficiency) and added to the activity database. Localisation information and date are extracted from the picture EXIF metadata. Any other information is erased at this step.

Once pictures are stored on the web interface, students can assign them to a specific quest and submit it for evaluation. The web application allows users to follow their progress in detail (which picture was submitted for which quest, what is the evaluation status, etc.) as well as the global progress of the other groups (the total number of collected points).

It is worth noting that in Belgium – where the web application was first used – the lockdown due to the COVID-19 pandemic still allowed citizens to go outside for some walk and exercise, although at a limited range. As such, the treasure hunt could still be performed by the students, either in their own garden or in neighbouring areas. However, not everyone lives in the countryside or close to a natural environment, or had the opportunity to leave their home during the lockdown. This is why we created a second module in the interface, the photo quiz, which allowed students to learn from photos contributed by other students, without having to submit their own photos.

### Photo quiz

The second module of the interface allows students to evaluate pictures submitted by other students (a modified version of peer evaluation). More precisely, in the photo quiz module, students are presented with pictures submitted by other groups and validated by the educators (see below “Expert evaluation). They have to assess whether the picture corresponds to its assigned quests. Their assessment is then compared to the assessment of the educators. If it matches, the students gain points that are added to their global group tally.

When performing this activity for the first time, it is necessary to have a sufficient amount of submitted (and corrected pictures). Without a database large enough, the activity loses some of its interest, as students might all review the same pictures.

### Expert evaluation

The third important module of the interface, central to the activity, is the expert evaluation. Each submitted picture needs to be manually assessed by the educators. Different feedback can be given for each submission, such as “correct”, “correct and nice picture”, “incorrect”, “not visible” (e.g. the object is not visible in the picture) or “out of rules” (e.g. picture of a houseplant, picture taken outside of the prescribed geographical zones). The interface was designed to easily navigate the different quests and quickly correct the submitted images.

### Technical aspects of the web application

The web application was created using the R Shiny framework, using the `shinydashboard` [7], `shiny` [8], `shinyWidgets` [9], `shinyBS` [10], `miniui` [11] packages for the user interface design. The data is stored in a SQLite database, hosted on the server. The database management is done using the `DBI` [12] and `RSQLite` [13] packages. Pictures are transformed and managed using the `magick` [14] package. EXIF information is extracted using the `exifr` [15] package. Data manipulation and visualisation is done using the `tidyverse` [16], `lubridate` [17], `cowplot` [18], `formattable` [19], `DT` [20], `plyr` [21], `leaflet` [22] packages. The text sentiment analysis was performed using the `rfeel` package [23].

In our exemple, the web application was deployed on the university server with the following specifications: Ubuntu 18.04.4 LTS x86-64, Linux kernel 4.15.0 x86-64, R 3.6.2 x86-64, Shiny server 1.5.12.933.

### Data accessibility

QuoVidi is an open source project, released under an APACHE licence [24]. Everyone is free to re-use and modify it, with attribution.

- Project website: http://www.quovidi.xyz
- Source code: https://github.com/QuoVidi/quovidi_public
- Script and data used for the manuscript: http://www.doi.org/10.5281/zenodo.3909033

## Results

### The web interface

The interface was created to be as much user-friendly as possible so that neither students nor staff need technical training. Because it is web based, it can be used on any platform, whatever the operating system. It scales on mobile devices as well, allowing users to store and submit pictures directly from the field (if they have an internet connection).

Figure 1 shows the different panels of the web interface. Figure 1A shows the “Store” panel, where students can store pictures, before submitting them for evaluation. This allows students from the same group to share and visualise their pictures. At this step, students can already assign a quest to the picture, which can be changed later on. They can also assign a geographic region, if this is required by the educators. A default region will be automatically proposed, based on the metadata of the picture.

**Figure 1:**
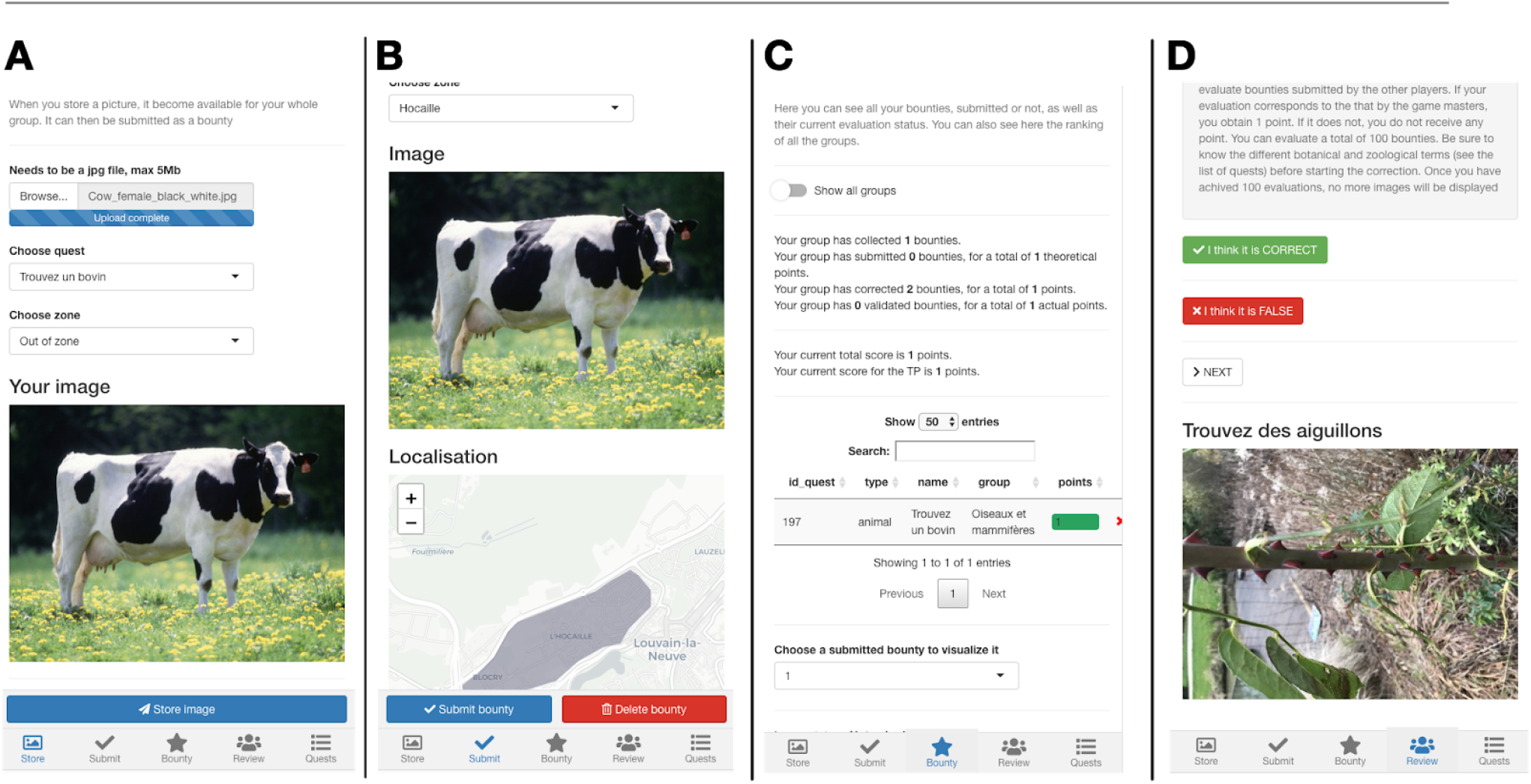
Overview of the different panels of the web interface. A. Store panel, where students can load the pictures taken in the field into the interface. B. Submit panel, where students can see all the stored pictures from their group and choose the ones to submit for evaluation. C. Bounty panel, where students can track the progress of their group and the others, as well as the expert evaluation of their submitted images. D. Review panel, where students can perform the photo quiz module.

Figure 1B shows the “Submit” panel. At this stage, students see all the pictures from their group. They can select a stored picture, assign it to a quest and submit it for evaluation. Groups can only submit one picture for each quest.

Figure 1C shows the “Bounty” panel, where students can visualise their progress. The panel presents an overview of the activity progress (for instance the total number of points or number of submitted quests). Students can also see the status of individual submissions, whether they are submitted or not as well as their validation status. In the same panel, students can also see the global scores of each group taking part in the activity. This adds a strong gamification aspect to the activity.

Figure 1D shows the “Quests” panel. In that panel students can navigate through the different quests proposed by the educators. They can sort them by groups, subgroups or rewards. In this panel, no explanation is given for the different quests. For instance if the quest is “Find an achene”, we do not define achene. This is done by design. We want students to look up the different biological terms by themselves. We do provide them with ressources to do so.

When an educator logs into the web application, the “Quests’’ panel becomes the “Admin” panel. In this panel, educators can follow the evolution of the activity (fig. 2A), change the activity parameters (fig. 2B) or correct the student submissions (fig 2C). Depending on the number of participating students and allowed submissions, the number of corrections can quickly become quite large. Therefore we designed the corrections interface to be fast and efficient. The educator first chooses one quest to correct. HeoShe will be presented with submissions for that quest only. The corrections are done in one click, on the appropriate feedback button. Previously validated submissions for this quest are presented on the side panel, to help maintain the consistency of the evaluations. The validated pictures are also a useful help for educators with a lesser expertise. Our experience shows that it takes, on average, 5-10 seconds to evaluate one submission.

**Figure 2:**
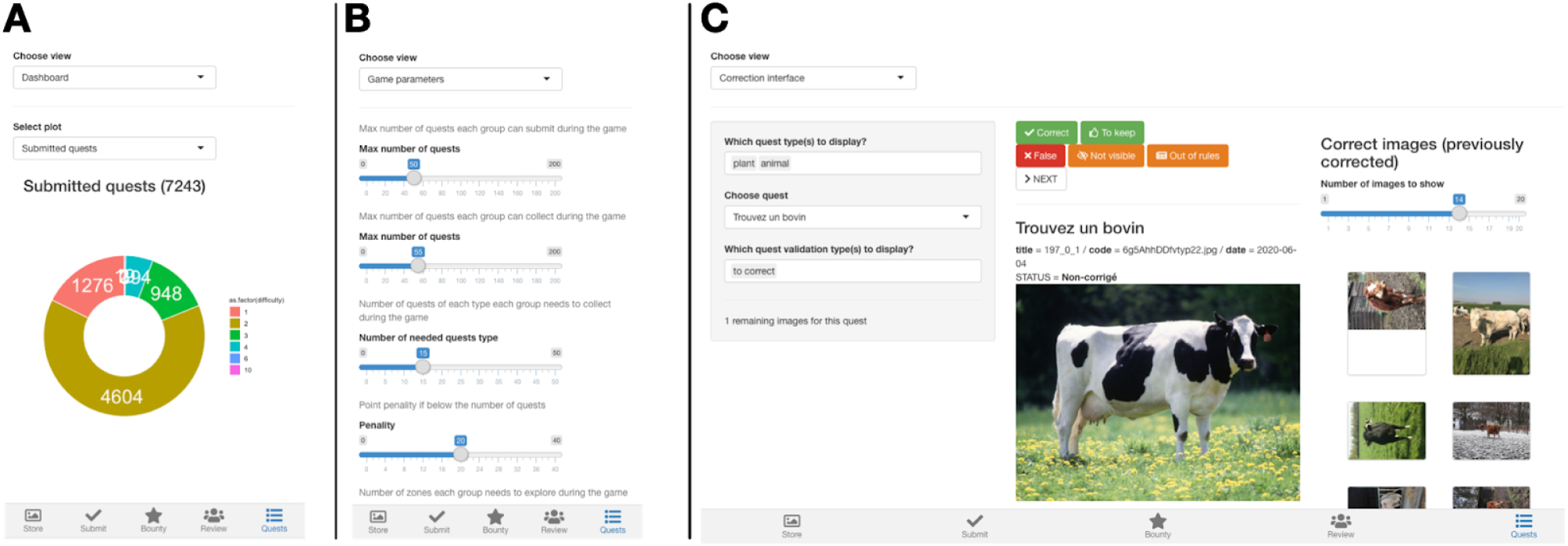
Overview of the different panels of the admin interface. A. Overview of the advancement of the game. For instance, educators can see how many pictures have been submitted and which proportion of these pictures has been evaluated. B. Game parameters. Educators can change the main game parameters directly through the web interface. C. Correction interface. The correction interface was designed to allow a quick and efficient correction process by the educators. The educator chooses a specific quest to evaluate, then simply clicks on the relevantfeed-back button. The right panel shows examples of previously validated images.

### The 2020 activity

In Spring 2020, we organised the activity with a rooster of 346 first year bachelor students from the Bioengineering Faculty of the UCLouvain (Belgium). Students were spread in 346 groups (it was therefore set up as an individual activity). Although students had to do the activity individually, we encouraged them to discuss the different quests and collect them together, as long as everyone took their own pictures. Each group was allowed to submit a maximum of 50 pictures. 285 quests were created, divided in 175 plant quests and 110 animal quests.

Specific restrictions were added to the game. A minimal number of animal and plant quests had to be collected by each group. Groups were also asked to collect pictures in different zones and biotopes (tab. 2) around the University campus, in Louvain-la-Neuve (Belgium).

**Table 2:** Description of the different zones defined for the 2020 activity. The total area of the game was 11.71 km2

| Zone name | Description | Area [km<sup>2</sup>] |
| --- | --- | --- |
| Bois de Lauzelle | Woody area | 4.85 |
| Hocaille | Urban area | 0.92 |
| Lac | Lake area | 0.18 |
| Bruyères | Urban area | 0.75 |
| Bois des Rêves | Woody area | 1.4 |
| Lauzelle-Centre | Urban area | 0.8 |
| Vieusart | Agricultural area | 1.81 |
| Biéreau-Baraque | Urban area with communal gardens | 1.0 |

The activity started on February 11. We had to pause the activity for 20 days at the beginning of the lockdown due to the COVID19 crisis. During that pause, we implemented the peer-evaluation in the web interface (it was not part of the interface initially). The activity resumed on the 3d of April and finished on the 15th of May. For the second phase of the activity, during the lockdown, all restrictions (quests groups and zones) were lifted as many students had returned to their home far away from the campus.

At the end of the activity, we sent an anonymous feedback form to the students and received 125 answers.

### Biological data collection

A total of 6543 pictures were submitted by students during the 2020 activity. Figure 3 shows the repartition of the submitted pictures by the students during the activity. Figure 3A & B show the difference before and after the lockdown imposed during the COVID-19 crisis.

**Figure 3:**
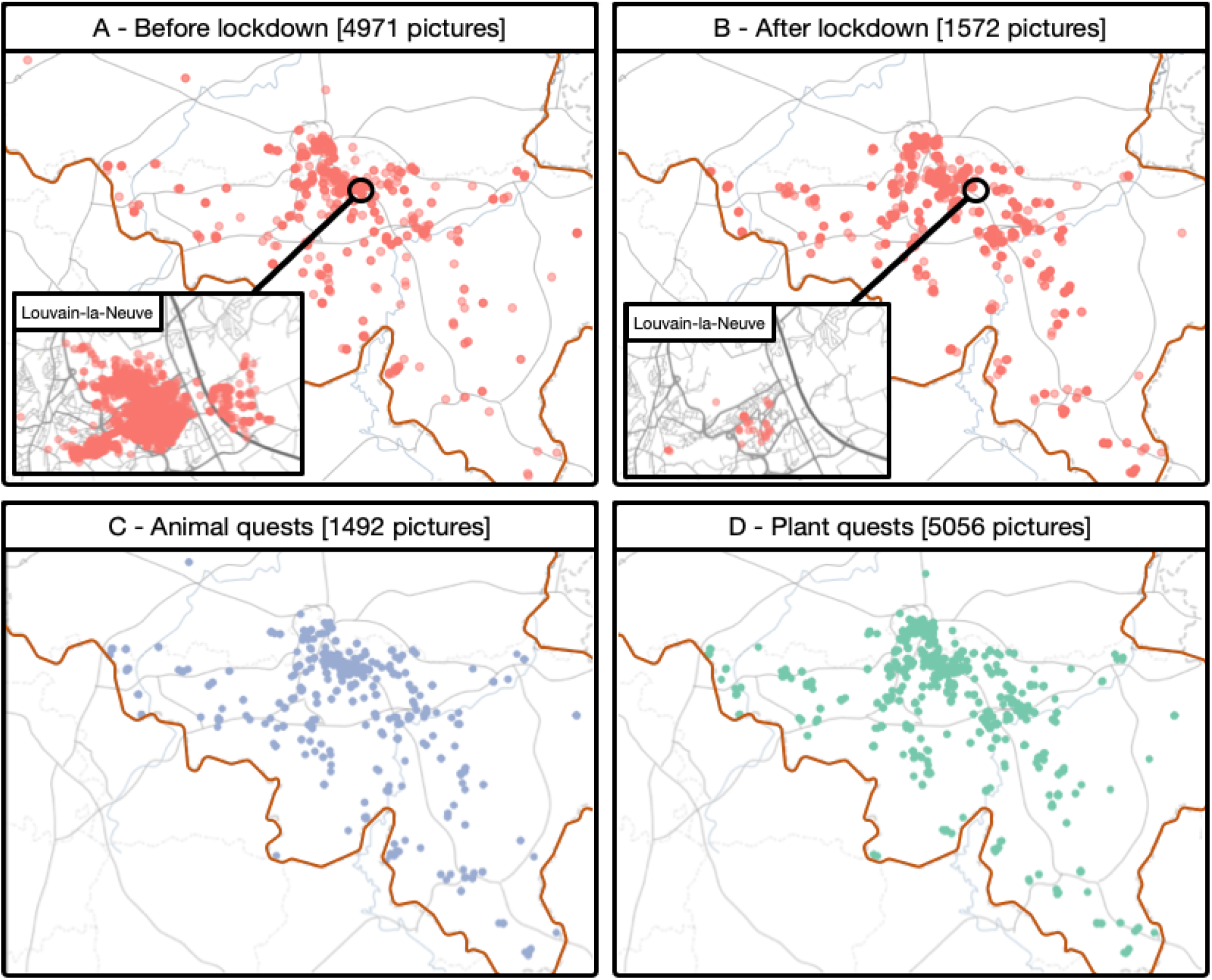
Overview of the data collected during the 2020 QuoVidi activity at the UCLouvain (Belgium). A. Pictures collected before the lockdown due to the COVID-19 crisis. B. Pictures collected after the lockdown. C. Animal quests collected during the whole activity. D. Plant quests collected during the whole activity. The belgian border is indicated in red

Before the lockdown, as we asked students to take pictures around the university, most of them were taken in Louvain-la-Neuve. During the lockdown, almost no pictures were taken in Louvain-la-Neuve, as students went back home. The lockdown reduced the number of collected pictures, but did not stop it. This is due to several reasons. At the beginning of the activity, we encouraged students to look for quests in groups, to foster peer-learning between them. This was not possible anymore during the lockdown. The collection of biological data was also influenced by the direct surroundings of the students. Students living in an urban area were potentially at a disadvantage compared to students in the countryside.

However, because we included the photo quiz module at the beginning of the lockdown, every student could continue the activity. Figure 4 shows, for every group, the proportion of points acquired either with the quests collection or the photo quiz. We can see that the dual system allowed students to choose different strategies, to adapt to their individual lockdown conditions.

**Figure 4:**
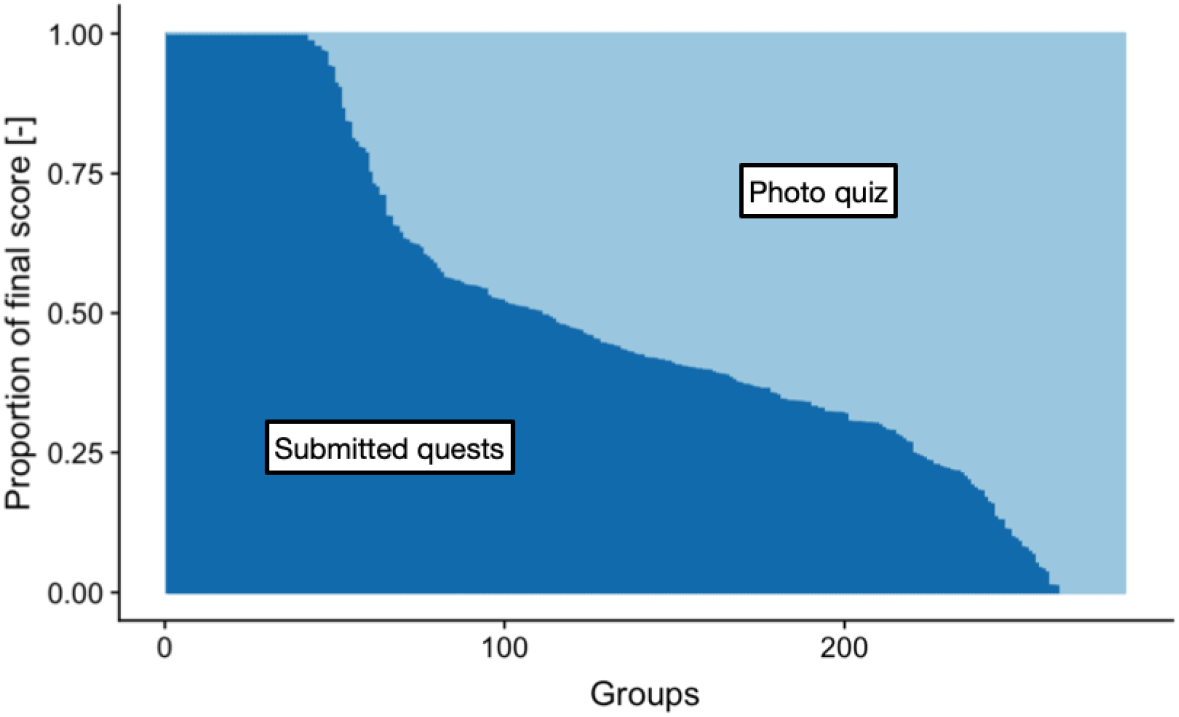
Proportion of submitted pictures and photo quiz points for each group.

We also observed a strong trend toward the collection of plant-related quests by the students (fig. 3C & D). This is probably due to the fact that, in an urban setup, plants are easier to find that animals. For an inexperienced naturalist, it is also probably easier to take pictures of plants than animals that have a tendency to escape. All the pictures can be viewed interactively at the address http://2020.quovidi.xyz

### Student accuracy

Overall, we observed a high correctness in the students picture submissions (fig 5A). For the treasure hunt and the picture collection, only 10% and 14% of the quests (for the animal and plant, respectively) were assessed as incorrect by ourselves. One reason for such a high accuracy from the students might be the high level of engagement required by the activity. They have to learn the vocabulary and discuss with other students, and go outside often in groups to find what they have identified as appropriate for a quest submission. In the ICAP framework [3], we believe this corresponds to the “Interactive learning” level, enabling the highest learning capabilities.

**Figure 5:**
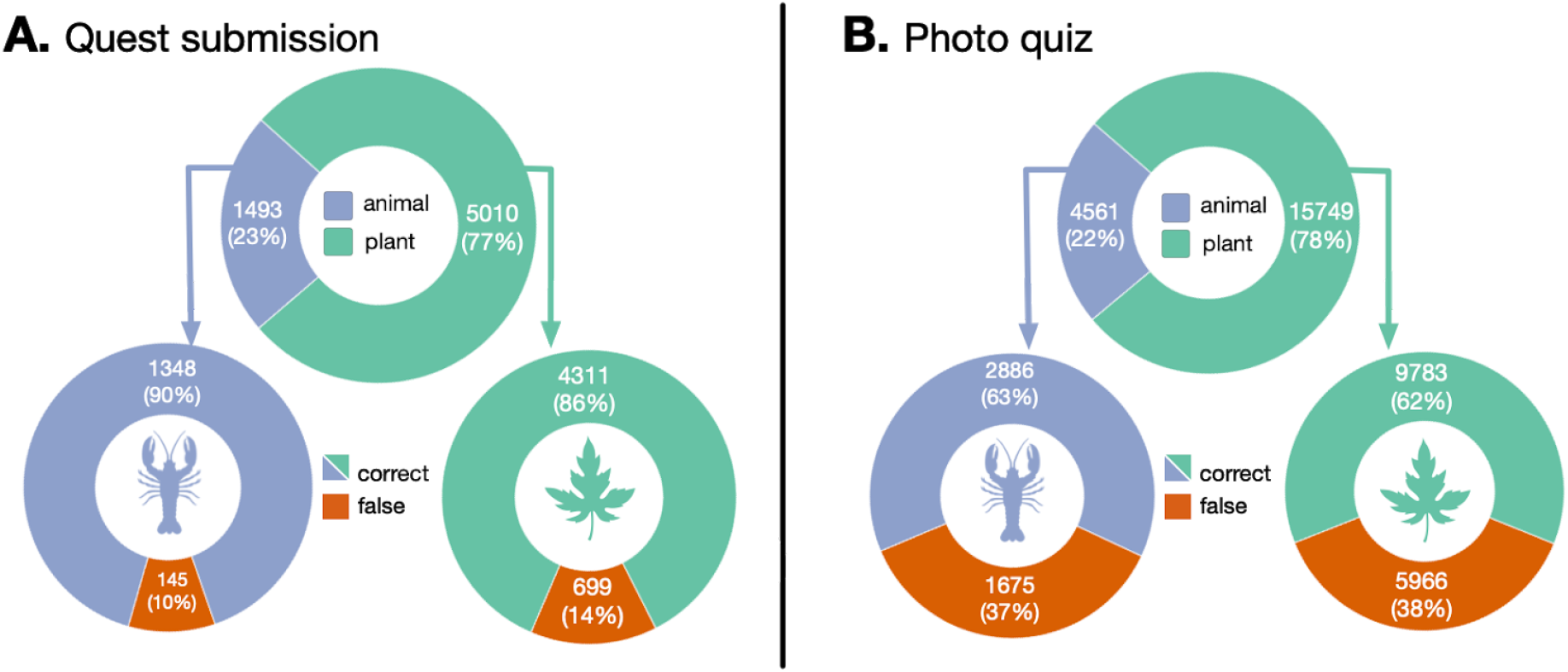
Performance of the students for the quest submission (A) and photo quiz (B) activities. In each panel, the top chart represents the proportion of plant and animal quests. The bottom panels represent, for each type, the proportion of correct and false submissions / corrections

Interestingly, we also observed a much lower accuracy for the photo quiz (fig. 5B). For that activity, 37% and 38% of the evaluations by the students (for the animal and plant, respectively) were incorrect. This can be due to several factors. First, contrary to the treasure hunt in itself, the evaluation activity requires a lesser level of engagement by the student. The activity is indeed “reduced” to click on a button in front of a computer screen. Second, depending on the quality of the picture to evaluate, said evaluation could be challenging. We tried to keep only good pictures for that activity, but the quality remained nonetheless variable.

### Students feedback

Overall, the activity was very well appreciated by the students. With a few exceptions, students like going outside to observe their surroundings and collect the quests. In a survey performed after the activity (fig. 6), 125 students reported to like the activity and have the feeling to have learned during it. Many students spontaneously expressed their enthusiasm for this activity (tab. 3).

**Figure 6:**
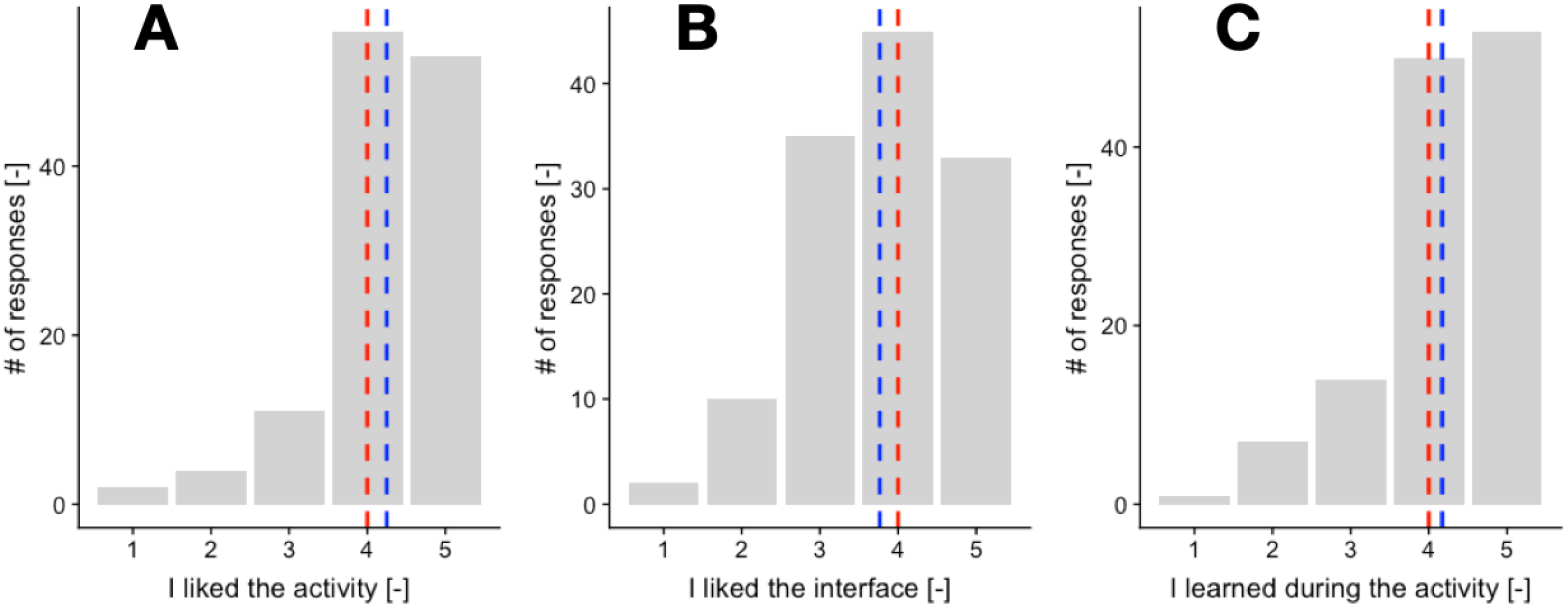
Feedback from the students. A. Global appreciation of the activity by the students. B. Appreciation of the web interface. C. Self assessment of learning during the activity. The numbers on the x-axis represent an increasing level of agreement with the statement presented, from strongly disagreed (1) to strongly agreed (5). Dashed red line represents the median while the dashed blue line represents the mean of the evaluation.

**Table 3:**
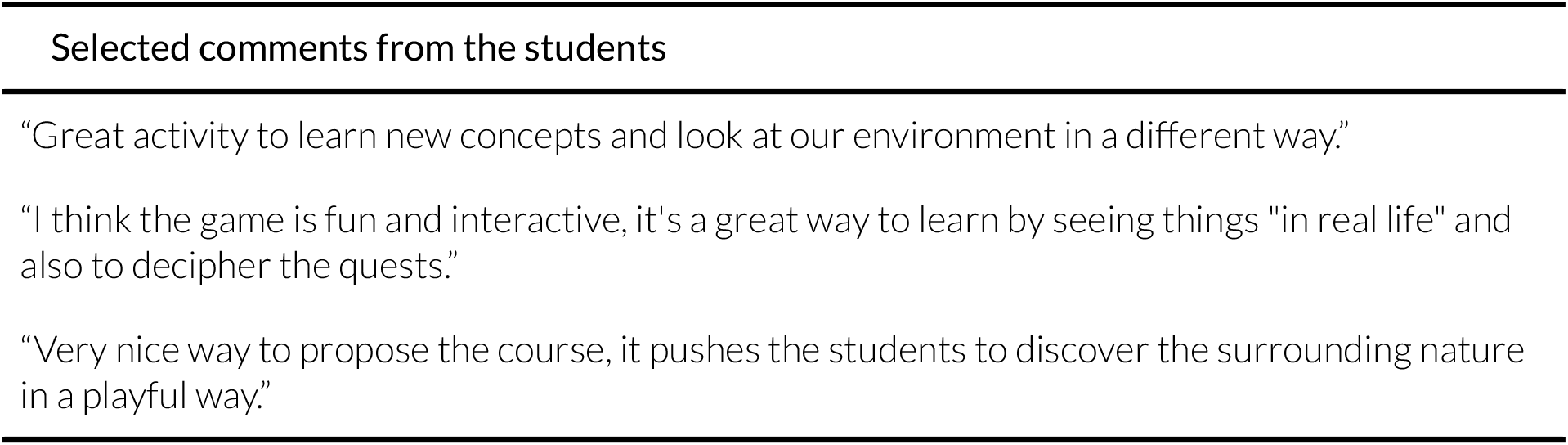
Selected comments from the students received with the feedback form.

## Discussions and perspectives

### Remote learning through a centralised game

The QuoVidi platform was created for several reasons. We wanted students to learn and know specific plant and animal vocabulary, but we did not want to just give them a list of words to be memorized and repeated. We also wanted them to explore and learn to observe their direct environment. We wanted to show them that you do not need to go to a tropical forest to be able to see a great diversity of plant and animal forms and species. We wanted to spark a strong interest in their surrounding natural world. Finally, we were also working with strong practical constraints. We needed to design an activity that was scalable for hundreds of students, without the need to increase the number of educators. This was possible, thanks to the current technologies (camera, mobile network and GPS localisation) available in almost every mobile phone.

With the creation of the web-platform for QuoVidi, we have met all those goals. The treasure hunt (and to a lesser extent the photo quiz) strongly motivates students to learn and remember the different technical terms used in the quests. Then they have to apply these new terms directly in the field. The gamification process (quests, score points, personal progress panel and scoreboard between all the groups) is also a strong incentive to engage in the activity.

The activity is also highly scalable. The number of participants is, from a technical point of view, only limited by the capacity of the server on which the platform is installed. The main limitation remains the expert correction step. As every single picture needs to be validated, the evaluation can quickly require a lot of time from the educator, even though we tried to make the process as efficient as possible. We hope in the future that the platform would benefit from advances in artificial intelligence algorithms to help correct the images (see below).

Finally, the activity is completely decentralised, which has been a great asset during the COVID-19 crisis. Students can collect quests at any time and place, making it easy to adapt to every individual situation. If they cannot go outside, or are not in a nature-rich environment, they can still participate in the activity via the peer evaluation module. From the educator point of view, all the management and corrections can be done from anywhere, as long as they have access to a computer and an internet connection. As such, the platform was a real asset during the lockdown period (13 March to 8 of June in Belgium), as it enabled us to continue the activity almost seamlessly.

### Reusing the image database

Similarly to citizen science projects, the use of our platform allows the collection of large numbers of geotagged, dated images of plant and animal structures. By helping create such a database over the years, the students are taking an active role in creating a valuable research ressource. This in itself is viewed by the students as a motivational element of the activity.

Such databases could be re-used in different ways. From an educational point of view, the images collected could be used to create a quiz to rehearse the vocabulary the following year. The student would therefore create their own teaching and rehearsal material. An example of a quiz created with the students pictures is visible here: http://quiz.quovidi.xyz.

From a research point of view, an ever growing database of annotated plant and animal pictures (describing either organ, species or groups), on a limited and well defined area would be a valuable resource. As each record of the database has been validated by an expert (the educators), such a database could be used in research projects.

Another interesting valuation of the database would be to reuse it to train deep learning recognition algorithms. Again, given the size and potential growth of the database, it will be an interesting resource to train machine learning models to recognise plant and animal structures. Such models could, in turn, be integrated into the platform to help with the correction.

### Collaborations between groups

So far, we use the QuoVidi framework within a single classroom (even if it was a very large one). Since the activity is entirely centralised online, we could imagine collaboration between remote classrooms. Students from different regions, countries or continents could participate in the same activity, hence increasing the degree of diversity of the observations.

### Expanding to new disciplines

Here we exemplified the use of our platform with a biological treasure hunt. Students were asked to find, in the field, plant and animal structures. However, due to its flexibility, the platform could be used to organise large scale treasure hunts in any context.

It could be used in architecture, design or geology classrooms, with quests related to different building structures, street art or rock, respectively. It could be used with children, with simplified quests, or with more advanced students, with more complex ones. In short, we expect the concept could be used in any context to deal with structures present in the “outside” world.

## Conclusions

We presented in this manuscript a new open-source web platform for the organisation for large tresor hunt, QuoVidi.

During the Spring 2020, in the midst of the COVID-19 crisis, we successfully used the QuoVidi platform with more than 300 students, and allowed the collection of more than 6000 geotagged plant and animal pictures. The decentralised nature of the platform enabled us to ensure a continuity in our teaching, despite the nation-wide lockdown.

We expect QuoVidi to be of interest for any teaching activity focused on the identification of real-world structures. QuoVidi is available at the address http://www.quovidi.xyz

## Authors contributions

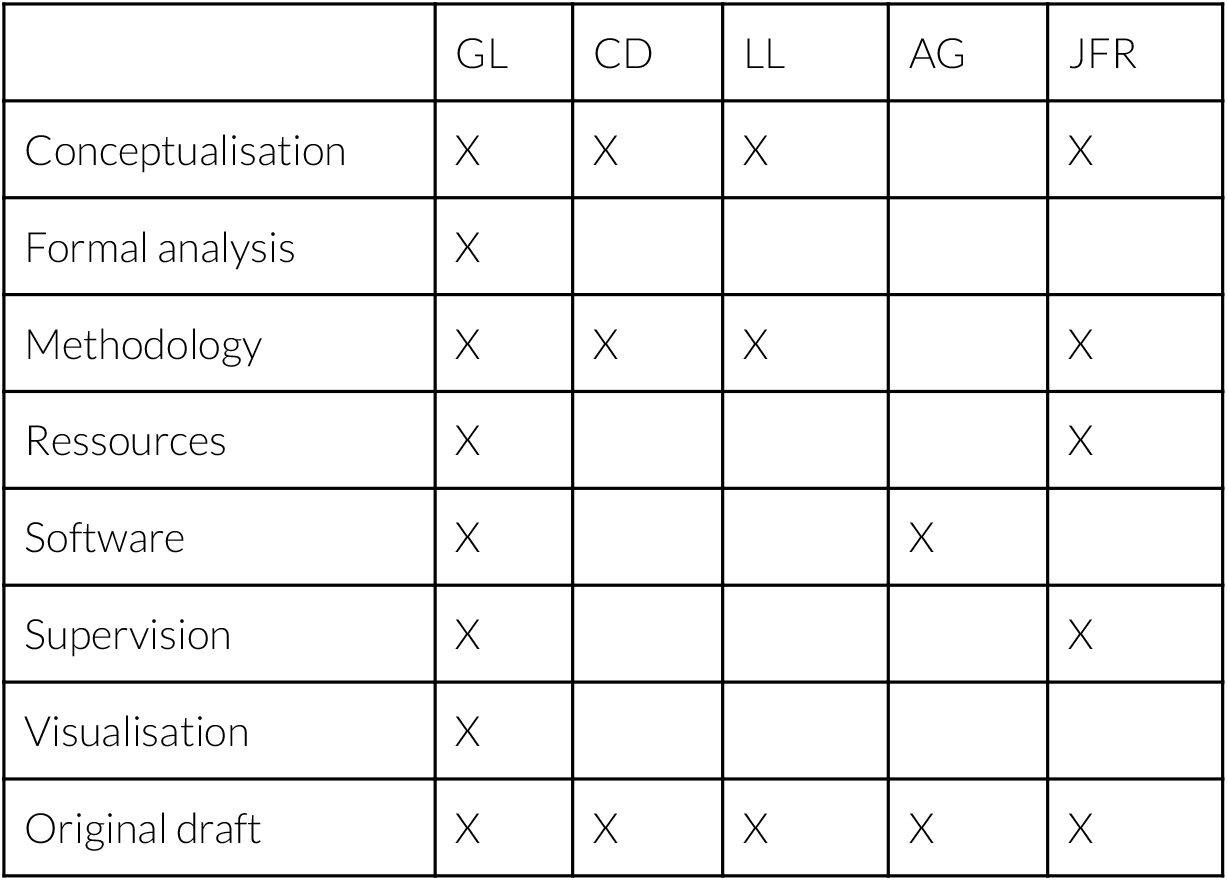

## Acknowledgments

QuoVidi, then called BioGO, was one of the laureates of the “Prix Wernaers pour la vulgarisation scientifique” in 2020.

## Notes

https://www.quovidi.xyz

### Competing Interest Statement

The authors have declared no competing interest.

